# Amyloid β Induces Hormetic-Like Effects Through Major Stress Pathways in a *C. elegans* Model of Alzheimer’s Disease

**DOI:** 10.1101/2024.05.07.593003

**Authors:** James D. Lichty, Adriana San Miguel

## Abstract

Amyloid β (Aβ) is a peptide known for its characteristic aggregates in Alzheimer’s Disease and its ability to induce a wide range of detrimental effects in various model systems. However, Aβ has also been shown to induce some beneficial effects, such as antimicrobial properties against pathogens. In this work, we explore the influence of Aβ in stress resistance in a *C. elegans* model of Alzheimer’s Disease. We found that *C. elegans* that express human Aβ exhibit increased resistance to heat and hypoxia, but not to oxidative stress. This beneficial effect of Aβ was driven from Aβ in neurons but not muscles, and the abundance of Aβ in neurons correlated with stress resistance levels. Transcriptomic analysis revealed that this selective stress resistance was mediated by the Heat Shock Protein (HSPs) family of genes. Furthermore, neuropeptide signaling was necessary for Aβ to induce stress resistance, suggesting neuroendocrine signaling plays a major role in activating organismal stress response pathways. These results highlight the potential beneficial role of Aβ in cellular function, as well as its complex effects on cellular and organismal physiology that must be considered when using *C. elegans* as a model for Alzheimer’s Disease.

## Introduction

Alzheimer’s disease (AD) is a progressive neurodegenerative disorder that causes loss of cognitive function in patients, typically starting at advanced age. AD is characterized by several features: the formation of amyloid β (Aβ) aggregates and micro-tubule associated protein tau (tau) tangles, immune activation, inflammation, oxidative stress, and neuron loss (Murke, 2009). In addition to forming plaques and aggregates, mutations in the Aβ peptide are associated with higher rates of AD, while Aβ expression also induces AD-like symptoms in cell and animal models (Alvarez et al., 2022, Blanchard et al., 2022, Yokoyama et al., 2022). These lines of evidence have made Aβ a priority target in AD research. Interestingly, some studies have also indicated that Aβ can potentially act as an antimicrobial peptide, suggesting a protective role for Aβ (Deepak et al., 2016, Pastore et al., 2020). Extensive work seeking to identify how Aβ becomes toxic in the brain has pointed to certain Aβ isoforms, small oligomer intermediates, and Aβ localization as key determinants in toxicity (Busch et al., 2022, Gallego et al., 2022, Rosenblum, 2002). Despite these discoveries, the role of Aβ in AD is still unclear, creating a need for well-characterized, robust models.

The nematode *C. elegans* has been previously used as a model for AD (Gallrein et al., 2022, Link et al., 2003, Wu et al., 2006). *C. elegans* boasts many benefits as a model for neurodegenerative diseases: it is transparent, allowing for *in vivo* imaging in live animals; has high homology with the human genome and proteome, including several stress-related pathways; and shares many neuron subtypes with humans (Corsi et al. 2015). Current models of AD in *C. elegans* focus on intracellular expression of human Aβ either pan-neuronally or pan-muscularly (Gallrein et al., 2022, Link et al., 2003, Wu et al., 2006). These models exhibit several pathological features such as severe paralysis, behavioral and motor defects, and reduced lifespan (Gallrein et al., 2022, Link et al., 2003, Wu et al., 2006). Aβ expression in worms has also been linked to loss of function and stability in mitochondria, leading to oxidative and metabolic stress (Teo et al. 2019). Conversely, worms expressing Aβ have also been shown to exhibit increased pathogen resistance, indicating a possible beneficial effect against environmental stressors (Deepak et al., 2016). Aβ shares some characteristics and behaviors with antimicrobial peptides, such as pathogen cell surface adhesion and pore formation, which may be the cause of this interaction (Deepak et al., 2016, Pastore et al., 2020). The relationships between Aβ and other environmental stressors have yet to be examined despite several stressors, like oxidative stress, being associated with higher rates of AD (Bisht et al., 2018, Ionescu-Tucker, Cotman, 2021). Investigating these interactions may improve our understanding of AD and how Aβ functions within it.

There is significant overlap between stress pathways in animals, including *C. elegans* and humans. The *C. elegans* gene *daf-16*, an ortholog for the human forkhead transcription factor (FOXO), has been highlighted for its ability to extend lifespan and increase stress resistance (Tissenbaum, 2018). Another *C. elegans* gene, *hsf-1*, is involved in the heat stress response and is an ortholog for the human HSF1 (Kyriakou et al., 2022). The *hif-1* gene (ortholog of human HIF1A) controls hypoxia resistance by regulating many hypoxia response genes, including *vhl-1* and *egl-9* (Powell-Coffman, 2010). The *skn-1* gene acts as a master regulator of the oxidative stress response in worms and its human orthologs, the Nrf/CNC proteins, are also regulators of oxidative stress (Blackwell et al., 2015). All four of these transcription factor pathways have important roles in aging and aging-related diseases in humans and have been implicated in AD (Calderwood, Murshid, 2017, Du, Zheng, 2021, Hassan, Chen, 2021, Yuan et al., 2018). Due to the substantial overlap between human and *C. elegans* stress response pathways, analysis of interactions between stress and Aβ in *C. elegans* can shed light on the effects of Aβ in humans.

In this work, we use several *C. elegans* models of AD to examine whether interactions exist between the effects of environmental stressors and Aβ expression and elucidate the underlying pathways behind any changes in Aβ-driven resistance to stress. We assessed the resistance of Aβ-expressing worms to oxidative stress, heat stress, and hypoxia. Our results indicate that Aβ expression selectively increases resistance to heat stress and hypoxia while reducing resistance to oxidative stress. When supplemented with an antioxidant, n-acetyl cysteine (NAC) and heat stressed, Aβ-induced stress resistance was unchanged, indicating that ROS (reactive oxygen species) does not play a role. Despite causing oxidative stress, the mechanisms by which Aβ increases heat and hypoxia resistance is not ROS-driven hormesis. Neuronal expression of Aβ affected stress resistance in a dose-dependent manner, while muscular levels of Aβ had no correlation with stress resistance. Using nCounter gene expression analysis, we identified several stress genes upregulated by Aβ in both unstressed and stressed worms. Several of these genes act downstream of the major stress response pathways modulated by *daf-16, hif-1, hsf-1*, and *skn-1*, suggesting Aβ activates these transcription factors either directly or by inducing cellular stress. Since stress response genes are mainly activated in the intestine, we sought to determine how neuronal Aβ was influencing other tissues to induce stress resistance. Using RNAi to knockdown two well-known genes required for neuropeptide signaling, *unc-31* and *egl-3* (Cornell et al., 2022, Salem et al., 2018), stress resistance returned to WT levels, pointing to neuropeptide signaling as the source of communication from neurons to other tissues for Aβ-induced stress resistance. These factors taken together highlight the complex effects of Aβ in cells and at the organismal level. Aβ expression has some beneficial effect for the animals in some contexts and thus results of studies of Aβ in *C. elegans* should consider possible interactions with environmental conditions. These results suggest that Aβ may play a substantial role in stress response, either through directly activating stress pathways or acting as a hormetic stressor.

## Materials and Methods

### Strains

**Table 1.** *C. elegans* strains used in this manuscript.

| Strain | Genotype | Source |
| --- | --- | --- |
| N2 | Wild type | From CGC |
| JKM2 | *Is [rgef-1p::Signalpeptide-Abeta(1-42)::hsp-3(IRES)::wrmScarlet-Abeta(1-42)::unc-54(3'UTR) + rps-0p::HygroR]* | Kirstein Lab<br>(Gallrein et al., 2021) |
| JKM3 | *Is [rgef-1p::wrmScarlet::unc-54(3'UTR) + rps-0p::HygroR]* | Kirstein Lab<br>(Gallrein et al.,<br>2021) |
| CL2355 | dvls50 [pCL45 (*snb-1::Abeta 1-42::3' UTR(long) + mtl-2::GFP*) I] | From CGC |
| CL4176 | dvls27 [*myo-3p::A-Beta (1-42)::let-851 3'UTR*) + *rol-6(su1006)*] X | From CGC |
| OH438 | otls117 [*unc-33p::GFP + unc-4(+)*] | From CGC |
| NL2099 | *rrf-3(pk1426)* II | From CGC |
| ASM35 | JKM2xNL2099 Cross | This work |

#### *C. elegans* Growth and Maintenance

Nematodes were grown and maintained at 20 °C on nematode growth medium (NGM) seeded with *E. coli* OP50 as a food source, unless otherwise specified, according to standard protocols (Wood, 1988). Age-synchronization for experiments was performed by washing plates with 1mL of a solution of M9 media supplemented with 0.01% v/v TX-100 (M9TX). Worms were allowed to settle, and supernatant was removed and replaced with 1mL of 1:2:1 mixture of bleach, 1M NaOH, and water. Once eggs were released, they were washed 3 times with 1mL M9TX, and transferred to fresh plates.

#### *C. elegans* Crossing

The JKM2 strain naturally produces a higher proportion of males than N2, so no extra procedures were needed to obtain males for the cross. To cross, young-adult NL2099 hermaphrodites were isolated onto a plate with JKM2 males in a 1:10 ratio, hermaphrodites to males. After several days, offspring were isolated and checked from homozygous passing of transgenes. The JKM2 transgene was checked using fluorescence microscopy. Once the JKM2 transgene was confirmed for 100 % transmittance, individuals were picked to fresh plates and allowed to grow. When sufficient offspring were on each plate (minimum 100 worms) genomic DNA was collected using standard methods and PCR was performed using primers flanking the *rrf-3(pk1426)* deletion site obtained from the Wormbase online resource. DNA showing the deletion was then processed through Sanger sequencing to confirm the *rrf-3(pk1426)* mutation.

#### Oxidative Stress Assay

Animals were age-synchronized by bleaching and cultured to the L4 stage at 20 °C. After another 24 hrs. of growth, approximately 30 worms were picked to a fresh plate containing 50mM paraquat (added during plate production after media was cooled to 50 °C) per cohort. These plates were transferred to a 20 °C incubator for 24 hrs. and then scored for survival by prodding with a platinum wire worm pick to check for movement. Animals that didn’t respond were marked as dead.

#### Heat Stress Assay

Animals were age-synchronized by bleaching and cultured to the L4 stage at 20 °C. After another 24 hrs. of growth, approximately 30 worms were picked to a fresh plate per cohort at room temperature (20 °C). These were transferred to an incubator set at 37 °C for the specified time (4 hrs. or 5.5 hrs.), before being returned to the 20 °C growth incubator for overnight recovery. For the 25 °C upshift experiments, these plates were instead first transferred to a 25 °C incubator for the specified time before the 37 °C stress. Animals were scored for survival by prodding with a platinum wire worm pick to check for movement. Animals that didn’t respond were marked as dead.

#### Hypoxia Assay

Animals were age-synchronized by bleaching and cultured to the L4 stage at 20 °C. After another 24 hrs. of growth, approximately 30 worms were picked to a fresh plate per cohort. These plates were put in a GasPak chamber with 3 satchels to lower the oxygen concentration to below 0.1 %. After 48 hrs. the chamber was opened, and the worms were allowed to recover overnight before scoring. Animals were scored for survival by prodding with a platinum wire worm pick to check for movement. Animals that didn’t respond were marked as dead.

#### RNA extraction and Gene Expression Profiling

Animals were age-synchronized by bleaching and cultured to the L4 stage at 20 °C. After another 24 hrs. of growth, worms were split into three treatment groups: unstressed, heat stressed, or hypoxia stressed. Unstressed worms were allowed to grow for another 24 hrs. before RNA was extracted. Heat stressed worms were stressed the next day at 37 °C for 2.5 hrs. Hypoxia stressed worms were stressed for 24 hrs. as described above. Following treatment, RNA was extracted according to the Direct-zol RNA Miniprep extraction kit (Cat# R2051). During the Tri Reagent step, a motorized pestle was used for 1 min to help break the worm cuticle. RNA samples were then provided to the UNC Respiratory TRACTS Core, who performed the Nanostring nCounter analysis. For each combination of strain and condition, three biological replicates were pooled together.

#### RNAi Plate Preparation and Suppression Assay

Bacteria and plates were prepared according to a modified protocol from Timmons et al. (Timmons et al. 2001). In summary, bacteria was cultured from the Ahringer RNAi Library (Kamath, Ahringer, 2023) in fresh LB media supplemented with 50 µg/mL ampicillin and 12.5 µg/mL tetracycline. After overnight growth (16-18 hrs.), the culture was centrifuged and resuspended in fresh LB media supplemented with 50 µg/mL ampicillin and 1mM Isopropyl-β-d-thiogalactopyranoside (IPTG) to OD_600_=0.5. After another 4 hrs. of growth, bacteria was seeded onto NGM plates supplemented with 50 µg/mL ampicillin and 1mM IPTG. Animals were age-synchronized by bleaching and cultured to the L4 stage at 20 °C on plates containing the relevant RNAi *E. coli* strain (control, *unc-31, egl-3*). After another 24hrs of growth, approximately 30 worms were picked to a fresh plate per cohort containing their respective RNAi *E. coli* strain. The heat stress assay was then performed as described above.

#### Statistical Analysis

Statistical analysis (ANOVA and t-test) was performed using MATLAB. P-values were considered significant when: p<0.05(*), p<0.01(**), and p<0.001(***).

## Results and Discussion

### Aβ induces selective stress resistance in *C. elegans*

To determine whether Aβ modulates responses to environmental stress, we analyzed the resistance of worms expressing Aβ to various stressors. We chose three strains for exposure: a wild-type N2 (WT), JKM2 (Aβ+), which co-expresses Aβ and Aβ tagged with wrmScarlet pan-neuronally, and JKM3 (Aβ-), which expresses wrmScarlet pan-neuronally (Gallrein et al. 2021). The Aβ+ strain expresses the tagged Aβ sub-stoichiometrically to limit wrmScarlet’s impact on Aβ aggregation, resulting in an aggregation pattern driven by Aβ (Gallrein et al. 2021). To assess worms’ resistance to oxidative stress, we exposed them to a 50 mM paraquat solution for 24 hrs. (Fig 1A). As expected, this toxic dose of paraquat (Gonzales-Moreno et al., 2023) resulted in reduced survival in the Aβ+ strain (37 %) as compared to the N2 wildtype (92 %) and the Aβ-strain (69 %) (Fig 1B). The Aβ+ strain exhibits elevated levels of internal oxidative stress from Aβ expression (Gallrein et al. 2021), possibly increasing their susceptibility to external oxidative stress and thus reduced resistance to paraquat. The Aβ-strain exhibited a less severe decrease in survival, possibly due to expression of neuronal wrmScarlet. Although the Aβ+ strain also expresses wrmScarlet, it does so at much lower levels than the Aβ-strain, and this effect does not completely account for the discrepancy in survival between the two strains.

**Fig. 1.**
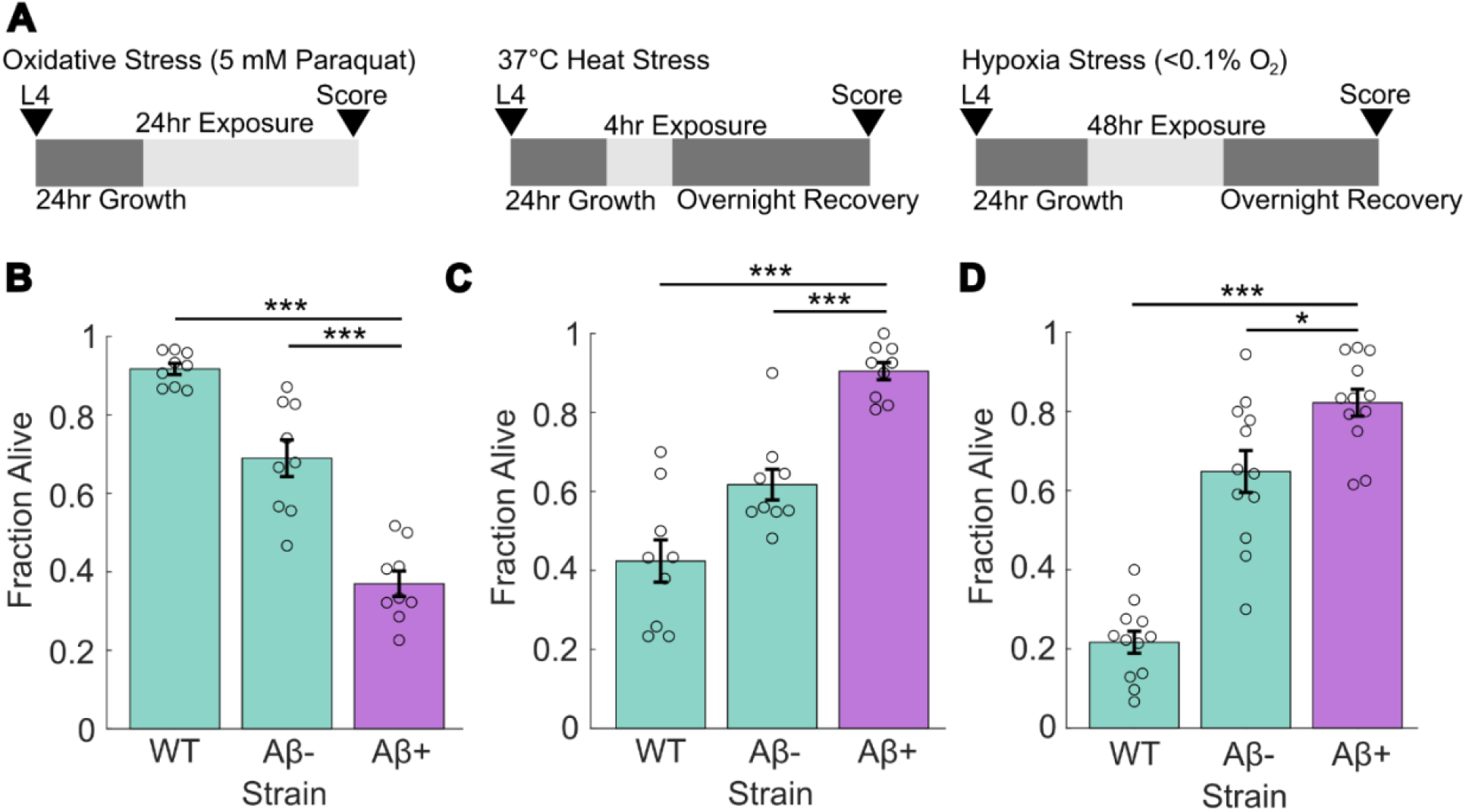
Aβ provides selective resistance to severe heat and hypoxic stress, but not paraquat-induced oxidative stress. (A) Overview of experimental setup. Animals were age-synchronized to L4 larvae and then allowed to grow for another 24 hrs. before exposure to stressors. (B) Survival rate after 24 hrs. exposure to 50mM paraquat stress. (C) Survival rate after 4 hrs. 37 °C heat stress exposure and overnight recovery at 20 °C. (D) Survival rate after 48 hrs. hypoxia (< 0.1 % O_2_) exposure and overnight recovery. WT is wild-type N2 Bristol, Aβ+ is JKM2, Aβ-is JKM3. N = 9 (12 for hypoxia stress) replicates per strain. Each dot represents a replicate of approx. 30 worms. Statistical analysis was performed using ANOVA. * is p-value < 0.05, ** is p-value < 0.01, *** is p-value < 0.005.

We next expanded our assay to include two additional common environmental stressors: heat stress and hypoxia. We exposed the worms to a severe heat stress of 37 °C for 4 hrs. and then allowed them to recover overnight at 20 °C (Fig 1A). Unexpectedly, the Aβ+ strain exhibited increased heat stress resistance, surviving at an average rate of 90%, compared to 42% and 62% in WT and Aβ-, respectively (Fig 1C). Similarly, when exposed to severe hypoxic conditions (<0.1 % O_2_) for 48 hrs. followed by overnight recovery (Fig 1A), the Aβ+ strain had increased hypoxia resistance, with an average survival rate of 82%, compared to 22% and 65% in WT and Aβ-, respectively (Fig 1D). We attributed this increased resistance to a possible hormesis-like effect. Notably, the Aβ+ strain does not display increased lifespan, which is often a hallmark of hormesis (Gallrein et al. 2021). The Aβ-control strain also demonstrated a higher tolerance to these two stressors when compared to WT, although not as high as the Aβ+ strain. This effect could stem from the wrmScarlet protein, as in the oxidative stress assay (Fig 1B). However, the substantial survival difference between the Aβ+ and the Aβ-indicate that wrmScarlet alone is not driving these differences. Aβ and wrmScarlet could induce similar protective effects, but the Aβ-strain lacks several other physiological deficits shown in the Aβ+ strain such as decreased lifespan, brood size, and locomotion (Gallrein et al. 2021).

### Foreign protein expression only partially accounts for increased stress resistance

As noted in our initial stress assays, wrmScarlet in the Aβ-strain induces some level of stress resistance. To test if foreign protein expression drives this effect, we sought to replicate the effect with another strain expressing GFP pan-neuronally, OH438 (referred to as nGFP) (Fig 3A) (Silveira et al., 2018). The nGFP strain exhibited increased heat stress resistance similar to the Aβ-strain, with an average survival rate of 53%. This result strengthens the idea that the hormesis-like effect stems at least partially from foreign protein expression in the neurons. Yet, the Aβ+ strain had significantly higher survival than the Aβ-strain (Fig 1), implying that foreign protein expression alone does not completely account for the effect and that Aβ induces protective effects against stress.

### Aβ-induced stress resistance is dependent on Aβ levels and localization

To validate that Aβ induces heat stress resistance, we tested the effects of increasing doses of Aβ in strain CL2355, referred to as the nAβ strain here. The nAβ strain expresses Aβ pan-neuronally upon upshift to 25 °C (Wu et al., 2006), and thus Aβ abundance is expected to be influenced by the time spent at 25 °C. We upshifted the nAβ strain to 25 °C for increasing amounts of time and then exposed it to heat stress (37 °C for 5.5 hrs.) followed by overnight recovery at 20 °C (Fig 2A-B) to test heat stress resistance. The nAβ strain exhibited increased heat stress resistance when compared to the WT strain, which increased with time at 25 °C. Notably, the 12 hr. timepoint showed the greatest difference in resistance between strains, with the nAβ strain exhibiting an average survival rate of 60% compared to the WT strain’s 33% average survival rate. We also noted that upshift to 25 °C produces an increase in heat stress resistance in the WT strain, which results in insignificant survival rate differences between strains at 24 and 48 hrs. The trend of 25 °C-induced heat stress resistance in wildtype animals is consistent with prior reports (Mendenhall et al., 2017, Servello, Apfeld, 2020). Despite this effect, our results suggest that Aβ expression induces an additional protective effect, since a significant difference in heat stress survival is observed between the two strains at 12 hrs.

**Fig. 2.**
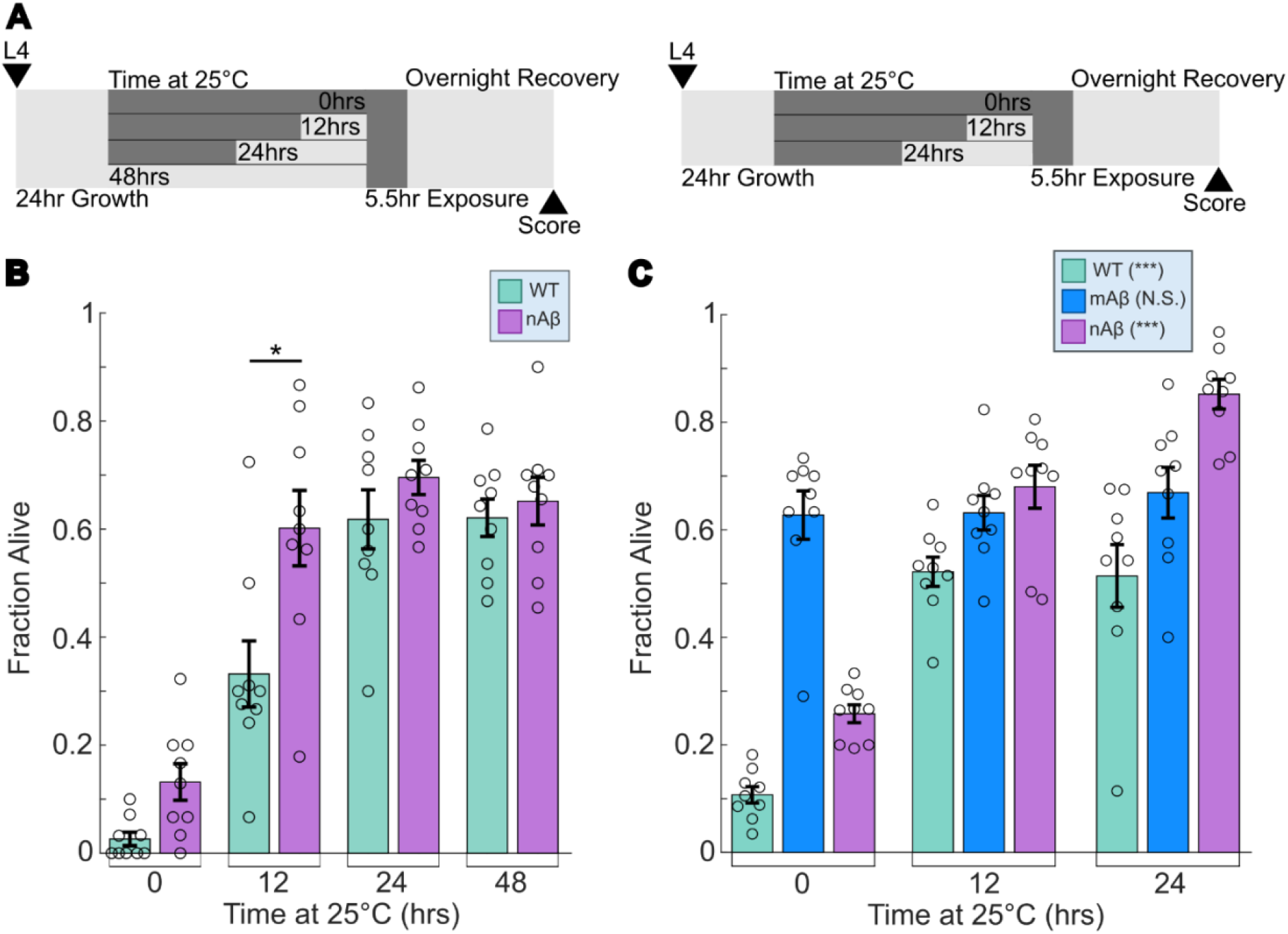
Aβ-induced heat stress resistance is dependent on Aβ expression levels and neuronal localization.(A) Overview of experimental setup. Temperature upshift to 25 °C was staggered such that stress exposure occurred at the same time for each condition. (B) Survival rate after 5.5 hr. 37 °C heat stress exposure and overnight recovery at 20 °C. Time at 25 °C correlates to levels of Aβ in the neurons in nAβ strain. (C) Survival rate after 5.5 hr. 37 °C heat stress exposure and overnight recovery at 20 °C. Time at 25 °C correlates to levels of Aβ in the neurons in nAβ strain and in muscles in mAβ strain. WT is wild-type N2 Bristol, nAβ is CL2355, mAβ is CL4176. N=9 replicates per condition. Each dot represents a replicate of approx. 30 worms. Statistical analysis was performed using two-sample t-test and two-way ANOVA. * is p-value < 0.05, ** is p-value < 0.01, *** is p-value < 0.005.

Since both the Aβ+ and the nAβ strains express Aβ in neurons, we next tested whether the increased stress resistance is specific to Aβ expression in this tissue. We expanded the heat stress assay to include strain CL4176 (referred to as mAβ) (Fig 2C). The mAβ strain also expresses Aβ upon upshift to 25 °C (Link et al., 2003), but in muscles instead of neurons. Rather than showing a dose-dependent heat stress resistance as we expected, the mAβ strain maintained a survival rate of approximately 64% regardless of the time spent at 25°C (Fig 2C). This result indicates that the resistance in the mAβ strain is not a result of Aβ levels. While the nAβ strain contains a fluorescence marker, the mAβ has a *rol-6(su1006)* marker that deforms the cuticle and results in a roller phenotype (Kramer, Johnson, 1993), which possibly explains this effect. In addition, the efficiency of Aβ production as a function of time at 25 °C likely differs in these two tissues. Since the mAβ strain’s resistance did not vary, while the nAβ strain’s resistance did vary and reached higher levels than the mAβ strain, we conclude that Aβ in neurons is a driver of heat stress resistance. It is unclear what aspect of the mAβ strain contributes to increased heat stress resistance, but previous work has shown that Aβ is undetectable when uninduced in this strain (Link et al., 2003), indicating that Aβ levels in muscles likely have little effect on resistance.

### Antioxidant exposure does not suppress the protective effect of Aβ

Since Aβ has been shown to induce oxidative stress in *C. elegans* (Drake et al., 2003), we hypothesized that the protective effect induced by Aβ could be the result of oxidative stress-driven hormesis (Cypser, Johnson, 2002, Ristow, Zarse, 2010). To determine if Aβ-induced oxidative stress drives increased stress resistance, we tested if the antioxidant NAC modulates the protective effect of Aβ. We exposed worms to 5 mM NAC for 24 hrs., a dosage that has been shown to reduce oxidative stress in *C. elegans* (Oh et al., 2015, Gusarov et al., 2021). After NAC exposure, animals were heat stressed at 37 °C for 4 hrs. and allowed to recover overnight. NAC had an insignificant effect on the Aβ+ and Aβ-strains but increased WT resistance (Fig 3B). NAC has been previously shown to induce some heat stress resistance (Oh et al., 2015, Gusarov et al., 2021), but it failed to significantly affect the Aβ+ strain. This suggests that Aβ-induced heat stress resistance is independent of any NAC-induced effects. Additionally, reducing Aβ-induced hormetic effects with NAC failed to produce a change, further indicating that Aβ modulates stress resistance in an oxidative stress-independent manner.

**Fig. 3.**
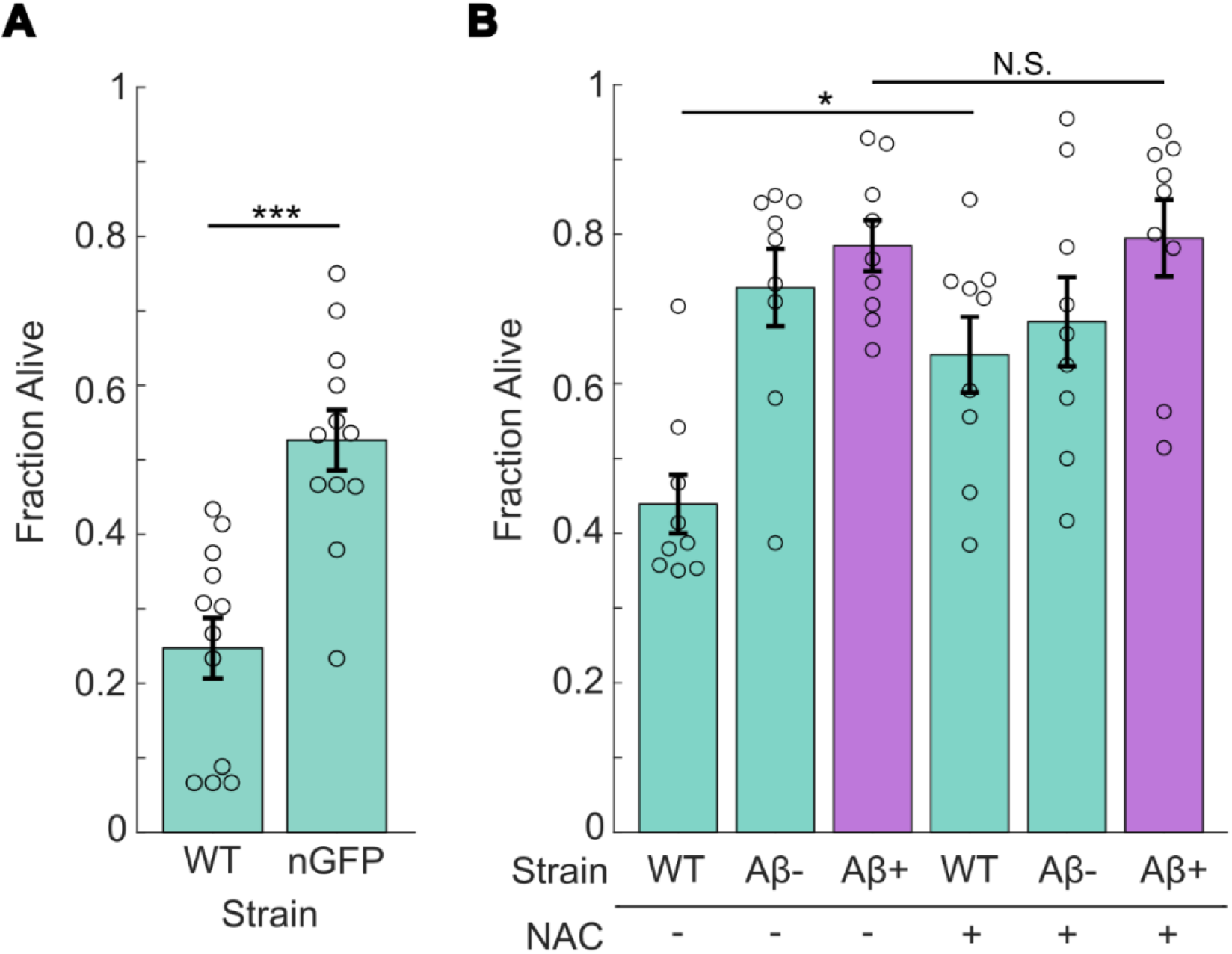
Heat stress resistance is partially replicated with other neuronally expressed proteins and is not eliminated with NAC treatment. (A) Survival rate after 4 hr. 37 °C heat stress exposure and overnight 20 °C recovery in pan-neuronal GFP strain. (B) Survival rate after 4 hr. 37 °C heat stress exposure and overnight recovery in worms grown on control and 5 mM NAC plates for 24 hrs. before heat stress exposure. WT is wild-type N2 Bristol, nGFP is OH438, Aβ+ is JKM2, Aβ-is JKM3. N=9 replicates per condition. Each dot represents a replicate of approx. 30 worms. Statistical analysis was performed using two-sample t-test and two-way ANOVA. * is p-value < 0.05, ** is p-value < 0.01, *** is p-value < 0.005.

### Aβ upregulates several stress response pathways

To elucidate how Aβ induces stress resistance, we probed the expression level of several stress resistance pathways using Nanostring nCounter analysis (Geiss et al., 2008). We targeted a predefined panel of targets that included several genes modulated by *daf-16, hsf-1, hif-1*, and *skn-1*, central regulators of the main stress response pathways. RNA samples from 3 biological replicates of each of the WT, Aβ+ and Aβ-strains were pooled for unstressed, heat stressed (2.5 hrs. at 37 °C), and hypoxic (<0.1% O_2_ for 24 hrs.) conditions (Fig 4A-C). This analysis revealed several stress resistance genes that were upregulated in the Aβ+ strain in both unstressed and stressed conditions, including several heat-shock proteins (HSPs). HSPs are associated with protein misfolding and molecular chaperones involved in the stress response, specifically heat stress, hypoxia, and oxidative stress (Kyriakou et al., 2022, Murshid et al., 2013). During stress, HSPs are typically upregulated in the neurons and intestine as a part of the stress response (Hodge et al., 2022). Aβ+ worms exhibited elevated levels of four HSPs under all conditions: *hsp-16*.*1, hsp-16*.*2, hsp-16*.*49*, and *hsp-70* (Fig 4A-C). This may indicate why Aβ induces more stress resistance than wrmScarlet, as the Aβ-strain only showed increased expression of these genes under hypoxia (Fig 4C). The HSPs are co-regulated by the *daf-16, hsf-1, hif-1*, and *skn-1* transcription factors (Hesp et al., 2015, Kyriakou et al., 2022, Park et al., 2009, Shamalnasab et al., 2017), making it unclear which pathway or pathways Aβ is inducing. Surprisingly, Aβ expression doesn’t upregulate any of the tested hypoxia or autophagy genes when compared to the controls, even under stressed conditions, suggesting the increased resistance is mediated mainly by the HSPs. Some of the main oxidative stress response genes, *sod-3* and *gst-4* (Oliveira et al., 2009, Wang et al., 2010), are slightly upregulated despite the Aβ+ strain exhibiting lowered oxidative stress resistance, which is likely explained by Aβ-induced oxidative stress. Interestingly, the *ttr-33* gene, another oxidative stress response gene responsible for paraquat and hydrogen peroxide resistance (Offenburger et al., 2018), is highly upregulated in unstressed worms, but this doesn’t translate to increased paraquat resistance when exposed (Fig 1B). Overall, this gene expression analysis suggests that Aβ induces stress resistance through upregulation of HSPs.

**Fig. 4.**
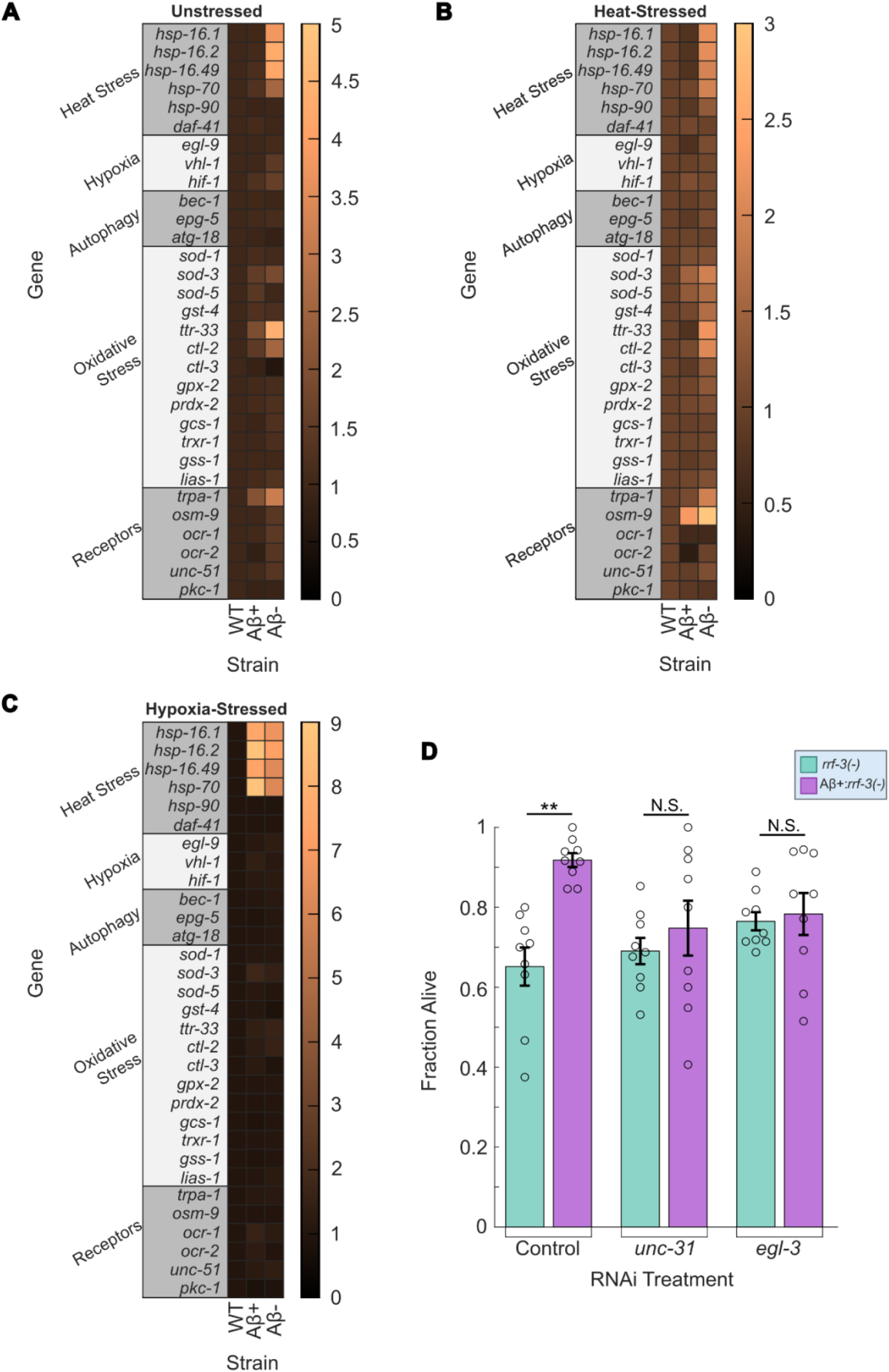
nCounter expression analysis indicates several resistance genes upregulated in both stressed and unstressed worms and RNAi suppression suggests resistance is communicated through neuropeptide signaling. (A-C) nCounter analysis on unstressed (A), heat stressed (B), and hypoxic stressed (C) worms. Genes are grouped by type and box color indicates expression changes. 3 biological replicates were pooled per sample. (D) Survival rate after 4 hr. 37 °C heat stress exposure in RNAi suppressed worms. RNAi control was HT115 with empty vector. WT is wild-type N2 Bristol, Aβ+ is JKM2, Aβ-is JKM3, rrf-3(-) is RNAi-sensitive strain NL2099, Aβ+rrf-3(-) is ASM35, a cross between JKM2 and NL2099. N=9 replicates per condition. Each dot represents a replicate of approx. 30 worms. Statistical analysis was performed using two-sample t-test and two-way ANOVA. * is p-value < 0.05, ** is p-value < 0.01, *** is p-value < 0.005.

### Aβ-induced stress resistance requires neuropeptide signaling

Stress resistance in *C. elegans* is a cell nonautonomous response mediated by communication between several tissues, most commonly the intestine and neurons (Hodge et al., 2022, Miller et al. 2020, Chen et al. 2024, Dutta et al., 2022). Stress leads to upregulation of transcription factors like *hif-1* in neurons, which can activate serotonin signaling and other pathways in a cell nonautonomous manner (Hodge et al., 2022, Kyriakou et al., 2022, Miller et al. 2020, Chen et al. 2024, Dutta et al., 2022). Neuroendocrine communication to the intestine through neuropeptide signaling can drive stress resistance through the activation of transcription factors in the intestine like *daf-16, skn-1*, and *hsf-1* (Hodge et al., 2022, Kyriakou et al., 2022, Miller et al. 2020, Chen et al. 2024, Dutta et al., 2022). The intestine can also act as a sensor for stress, such as detecting dietary restriction through food intake and *daf-16* activation, oxidative stress through *skn-1* activation, and heat stress through *hsf-1* activation (Hodge et al., 2022, Kyriakou et al., 2022, Miller et al. 2020, Chen et al. 2024, Dutta et al., 2022). The intestine can then communicate these signals back to the neurons to trigger an organism-level stress response (Hodge et al., 2022, Kyriakou et al., 2022, Miller et al. 2020, Chen et al. 2024, Dutta et al., 2022). Therefore, we next focused on whether neuronal Aβ could activate stress resistance pathways in the intestine through neuroendocrine communication. It has been previously shown that Aβ can spread from neurons to other tissues in aged *C. elegans* (Gallrein et al. 2021). In our experiments, we evaluated young worms, before the age where Aβ dispersal has been identified (Gallrein et al. 2021), thus ensuring Aβ is restricted to neurons. To understand how neuronal Aβ influences organism-level stress resistance, we assessed whether disrupting neuropeptide signaling between the neurons and other tissues had an effect on stress resistance. To knockdown neuropeptide signaling, we used RNAi by feeding (Timmons et al. 2001), targeting two key neuropeptide signaling genes, *unc-31* and *egl-3. unc-31* regulates neuropeptide release from dense core vesicles (Cornell et al., 2022) and *egl-3* is necessary for neuropeptide maturation (Salem et al., 2018); knocking out either severely limits neuropeptide signaling. To ensure silencing was effective in neurons, we used an RNAi sensitive strain, NL2099 (*rrf-3*(-)) (Simmer et al., 2002), as our control. This strain was crossed with the Aβ+ strain to generate ASM35 (Aβ+;*rrf-3*(-)). Upon knockdown of either gene (Fig 4D), Aβ-induced stress resistance in Aβ+;*rrf-3*(-) worms dropped to similar levels as *rrf-3*(-). This result suggests that Aβ in neurons upregulates stress resistance pathways in other tissues through neuropeptide signaling, thus driving increased organismal resistance to stress. It is unclear if Aβ is directly initiating neuropeptide signaling, or if Aβ induces neuronal stress that may be responsible for the activation of the stress response at the organismal level.

## Conclusion

In summary, this study found that Aβ induces resistance to heat and hypoxic stress but not oxidative stress by activating key stress response genes. Several members of the HSP family of genes were induced in the Aβ-expressing strain under both unstressed and stressed conditions. The location and levels of Aβ have significant effects on the levels of stress resistance induced, with muscular Aβ expression levels having no correlation with resistance. The stress resistance may be partially explained by neuronal expression of foreign protein, but Aβ is able to provide higher stress resistance than either of the two other proteins tested. It is unclear if protein aggregation plays a role in this protective effect, as both Aβ and wrmScarlet-tagged Aβ aggregate (Gallrein et al., 2021), but aggregation levels were not assessed in this work. Gene expression analysis indicates that Aβ activates some combination of the *daf-16, hsf-1, hif-1*, and *skn-1* pathways to induce stress resistance, but further work is needed to determine if all four pathways participate in this effect. The HSP family of genes falls under numerous transcriptional regulators, and expanding the breadth of genes covered by this kind of screen may provide more insight into which pathways are activated. RNAi knockdown of *unc-31* and *egl-3* shows that to induce stress resistance, Aβ activates neuropeptide signaling to communicate with other tissues. More work is needed to examine exactly how Aβ activates neuropeptide signaling, whether that be directly or indirectly by inducing cellular stress. This study highlights the need to more fully characterize the effect of Aβ on *C. elegans*, and to consider its complex organismal effects when using it as a model for AD. If the increased stress resistance is not consistent across species, there may be more limitations of *C. elegans* as a model for AD and more factors to consider before its use as such. Knowing these limitations and interactions will make interpretations of results more meaningful. The results of this work also imply that Aβ may have beneficial effects for the host, outside of AD. The role of Aβ in healthy humans is still quite unclear, but several theories regarding its function include pathogen defense, injury recovery, and regulation synaptic function (Brothers et al., 2018). Inducing stress resistance pathways and acting as an early signal for stress may be another role it plays in human biology. It will be crucial to further characterize the role of Aβ in *C. elegans* and other model organisms to better understand the mechanisms by which Aβ induces physiological changes in AD and other human diseases.

## Acknowledgements

Some strains were provided by the CGC, which is funded by NIH Office of Research Infrastructure Programs (P40 OD010440). We thank Janine Kirstein for sharing strains JKM2 and JKM3 and for useful insights on strain culture and handling. We also thank Wormbase and our funding sources DOD (W81XWH-18-1-0701) NIH (R00AG046911) and NSF (2039226).

